# DeepKin: Predicting relatedness from low-coverage genomes and paleogenomes with convolutional neural networks

**DOI:** 10.1101/2024.08.08.607159

**Authors:** Merve N. Güler, Ardan Yılmaz, Büşra Katırcıoğlu, Sarp Kantar, Tara Ekin Ünver, Kıvılcım Başak Vural, N. Ezgi Altınışık, Emre Akbaş, Mehmet Somel

## Abstract

*DeepKin* is a novel tool designed to predict relatedness from genomic data using convolutional neural networks (CNNs). Traditional methods for estimating relatedness often struggle when genomic data is limited, as with paleogenomes and degraded forensic samples. *DeepKin* addresses this challenge by leveraging two CNN models trained on simulated genomic data to classify relatedness up to the third-degree and to identify parent-offspring and sibling pairs. Our benchmarking shows *DeepKin* performs comparably or better than the widely used tool *READv2.* We validated *DeepKin* on empirical paleogenomes from two paleological sites, demonstrating its robustness and adaptability across different genetic backgrounds, with accuracy >90% above 10K shared SNPs. By capturing information across genomic segments, *DeepKin* offers a new methodological path for relatedness estimation in settings with highly degraded samples, with applications in ancient DNA, as well as forensic and conservation genetics.

## Introduction

Relatedness estimation using genetic data has uses in a wide range of fields of life sciences, from evolutionary and conservation genetics to anthropology and forensics. Close relatives also need to be identified in population genetic analyses and genome-wide association studies, where they represent nuisance factors to be removed. Although the classical tools for close kin identification have been microsatellites (STRs), a growing number of studies today perform relatedness estimation using genome-wide SNP data instead (*1*). When haplotype-level data and an appropriate background population sample for inferring haplotype frequencies are available, identifying close relatedness is a relatively simple problem (*2*). However, high-quality haplotype-level data is frequently unavailable in various fields of biological research, including ancient DNA, conservation biology (where samples may be derived from environmentally-collected material such as faeces), or forensic genetics (where biological material may also be highly degraded). Due to the degraded nature of DNA in these samples, an individual may be represented by only dozens of thousands of SNPs and only a single allele per SNP (i.e., in pseudohaploid form) due to low coverage. A pair of individuals being identity-by-state (IBS) per SNP contains only minute information on their relatedness, which renders estimating kinship with limited SNP numbers a significant challenge.

To date, different statistical tools have been developed for estimating relatedness using such sparse genetic data: *READv2* (*3*), *TKGWV2* (*4*), *BREADR* (*5*), GRUPS-rs (*6*), *ngsRelateV2* (*7*), *lcMLkin* (*8*), *KIN* (*9*), among others. These are being used particularly intensely in the rapidly growing field of human paleogenomics to study past social structures (*10*). These tools differ in whether they use population allele frequencies for normalising IBS to estimate relatedness or average IBS (or, inversely, mismatch) signals in a population sample. The methods also vary in the use of genotype likelihoods versus genotype calls. The most commonly used approach (*11*) in the ancient DNA literature to date is implemented by the software *READ*, which calculates genome-wide mismatch rates between a pair using genotype calls and normalises this value using the average mismatch rate in a population sample (*12*, *13*). Its most recent version, *READv2* (*3*), also distinguishes between first-degree related pairs, i.e., parent-offspring and siblings, using variation in IBS signals across the genome.

In a recent benchmarking study using simulated genomes, we studied the performances of four such statistical tools designed to estimate close kinship with limited data: *READ* (*13*), *ngsRelateV2* (*7*), *lcMLkin* (*8*), and *KIN* (*9*). We found that these methods have sufficient power (F1>0.90) to identify the correct kinship up to third-degree relatedness when two genomes share >20K SNPs (*11*). However, accuracy dropped considerably at lower SNP numbers, especially when distinguishing second-degree from third-degree relatives. A benchmarking study using empirical ancient DNA data likewise reported limited accuracy at comparably low genomic coverages (*14*). These constraints in accuracy could be partly attributable to random noise but also to the inherent randomness in meiosis (e.g., the fact that siblings may share more or less than half of their DNA sequences).

The threshold of 20K genome-wide SNPs shared between a pair may not always be achievable, especially in settings where samples are highly degraded or budgetary constraints hinder data generation. This calls for novel approaches for kinship estimation to apply to the lowest preserved samples. One possible strategy could involve using information across linked genomic segments instead of treating each SNP independently. In fact, the tool *KIN* uses a hidden Markov Model (HMM) precisely for this purpose, estimating IBD across genomic segments. According to our benchmarking study, however, the accuracy of *KIN* does not systematically exceed its simple alternative, *READ* (*11*).

Deep learning tools have recently shown exceptional performance in solving classification problems. A particularly attractive set of tools are convolutional neural networks (CNNs or ConvNets) (*15*), which consist of linear convolution operations, nonlinear activation functions, pooling layers, and fully-connected classification layers. CNNs are highly adept at handling grid-like data structures, such as images, as well as summaries of genomic data, which can likewise be stored in data arrays and processed like images. Indeed, CNNs have been successfully applied to genomic summary statistics for categorisation tasks in various population genetic applications, such as detecting positive selection or demographic inference, as reviewed by Korfmann et al. (*16*).

Here, we present *DeepKin*, composed of two CNN models trained on simulated genomic data from known pedigrees to classify pairs into relatedness groups using limited genomic data. *DeepKin* performs comparably and frequently better than *READv2* on simulated training data and two empirical datasets.

## Methods

See **Figure 1** for an overview of the basic workflow of this section.

**Figure 1:**
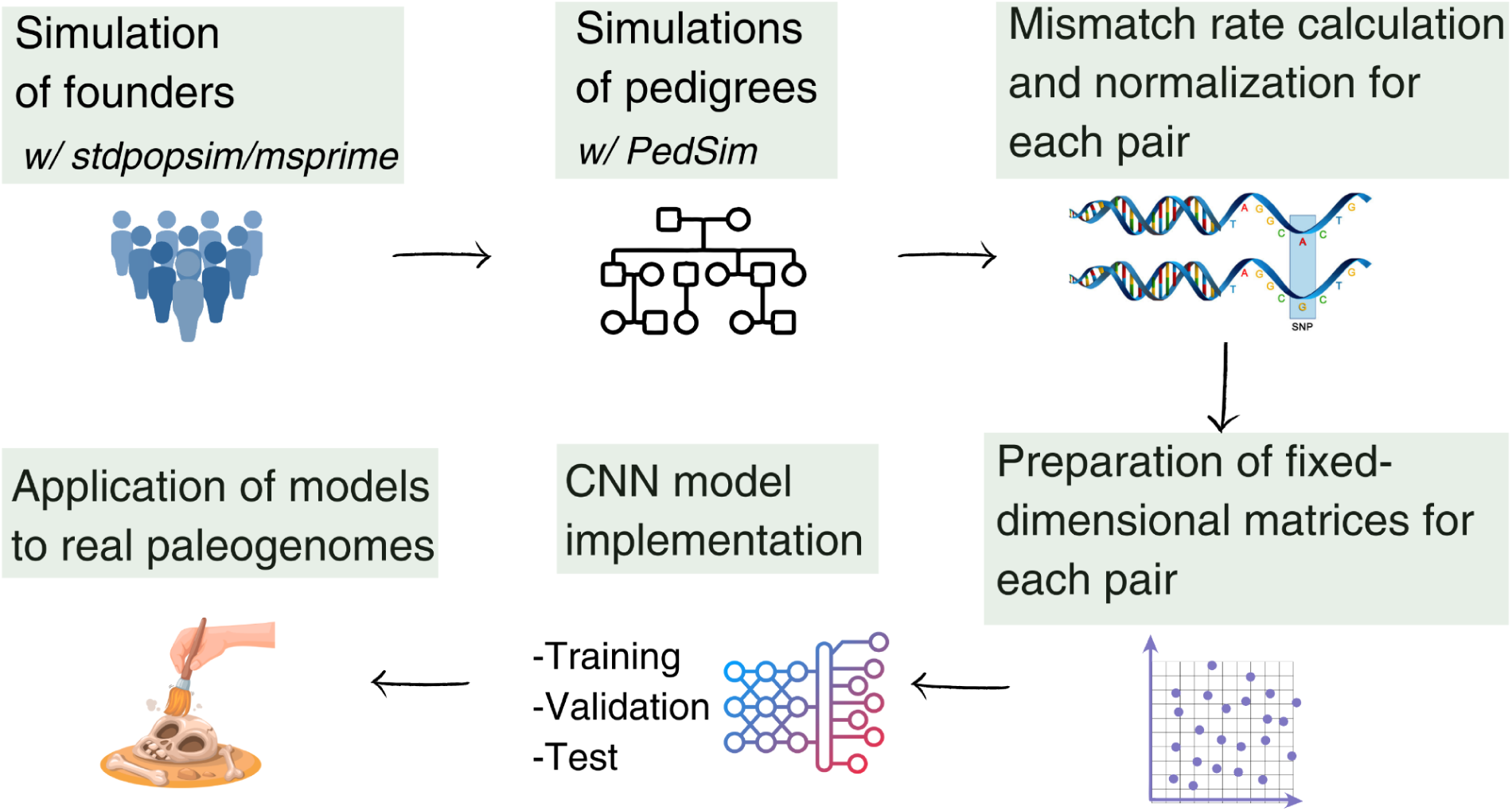
Overview of the workflow. The workflow begins with the simulation of founder genomes, followed by the simulation of pedigrees using the founders. Next, mismatch rates are calculated and normalised for each pair, yielding estimates of the relatedness coefficient (*r*) across genomic windows. Fixed-dimensional *r* matrices are then prepared for each pair and used as input for the CNN model implementation. Two configurations of CNN models are employed: Model-A (window size 200 SNPs, sliding window 50 SNPs) and Model-B (window size 500 SNPs, non-overlapping windows). These models undergo training, validation, and testing. Finally, the trained models are applied to empirical paleogenomes to predict relatedness.

### Simulation of founders

We created genomes for 250 founder individuals to be used as input in pedigree simulations based on realistic parameters, including chromosome size, mutation and recombination rates, and demographic history. For this, we employed the *msprime* engine (*17*, *18*) in the “HomSap” mode of the *stdpopsim* (*19*, *20*) library. The empirical recombination map was chosen as “HapMapII-GRCh37*”* (*21*) with the “-g” option in *msprime*. As a demographic model, we used an available multi-population demographic history model of ancient Eurasia, called from the *stdpopsim* library with the “-d AncientEurasia-9K19 0 500” option. From the model, we sampled genomes representing the Linearbandkeramik (LBK) group, a Neolithic farming population in central Europe with Anatolian origins (*22*). Our choice of the LBK was motivated by our goal to create a tool directly implemented to assess relatedness among West Eurasian prehistoric paleogenomes, a recent research focus in the field (*23–26*). We thus produced 500 haploid genomes. Subsequently, we used the *tskit* (*18*) library’s “vcf” command with the “--ploidy 2” option to convert the succinct tree sequence haploid output from *stdpopsim* into VCF format with diploid genotypes. Using a custom bash script, we selected 200,000 random SNP positions from the data. These positions were then utilised to extract reference bases from the human reference genome (hs37d5) with the “getfasta” command from *BEDtools* (*27*) (v2.27.1). We estimated the transition/transversion ratio from the 1000 Genomes Project (*28*) Dataset v3 for the Tuscany (TSI) population to assign alternative alleles to the reference positions. Using this information, we stochastically generated alternative alleles for each position in our dataset with a custom *R* script. We randomly assigned sex to produce 125 female and 125 male founders.

### Simulation of pedigrees

We simulated pedigrees, presented in **Table 1**, that included all types of relatedness up to the third degree. These relationships encompassed identical pairs, first-degree (parent-offspring, siblings), second-degree (half-siblings, avuncular, grandparent-grandchild), and third-degree (first cousins, great-avuncular, great-grandparent-great-grandchild) pairs, as well as unrelated individuals. We utilised the *Ped-sim* (*29*) pedigree simulator to simulate pairs in all relatedness categories except for the identical and unrelated pair categories. We provided the software with the phased VCF file, free of missing sites, containing simulated founder individuals using the “-i” option. We also specified the founder sexes of the input VCF samples with the “--sexes” option. We interpolated the physical positions in the VCF file to generate a genetic map file using the “filter_vcf.py” script from the *adna_tools* (*30*) *Python* package. This map file was used in the pedigree simulation step with the “-m” option. We applied the crossover interference model (*31*) provided by the *Ped-sim* software using the “--intf” option.

**Table 1:** The summary of the simulated dataset. The table summarises the details of the simulated dataset, including various relatedness types, the number of unique pairs, the number of downsampled versions, and the total number of pairs. The dataset is divided into two downsampling sets. Set 1 includes versions with overlapping SNP counts of 1,000, 1,500, 2,000, 2,500, 3,500, 4,000, 4,500, 6,000, 7,000, 8,000, 12,500, 15,000, 20,000, 30,000, 35,000, 40,000, and 100,000. Set 2 includes versions with overlapping SNP counts of 3,000, 5,000, 10,000, 25,000, and 50,000. The table also provides the distribution of pairs into training, validation, and test sets for each relatedness type and downsampling set. The "Labelled as" column provides information about the labels used while training the model. Notably, the total number of pairs across all relatedness types and downsampling sets is 230,242 for the training set, 57,561 for the validation set, and 17,338 for the test set.

| Relatedness type | Number of unique pairs | Number of downsampled versions | Total pair | Labelled as | Downsampling set | Training set | Validation set | Test set |
| --- | --- | --- | --- | --- | --- | --- | --- | --- |
| Identical | 1254 | 22 | 27588 | Identical | 1 | 17,054 | 4,264 | 0 |
|  |  |  |  |  | 2 | 3,762 | 941 | 1,568 |
| Parent-offspring | 1200 | 22 | 26400 | Parent-offspring | 1 | 16,320 | 4,080 | 0 |
|  |  |  |  |  | 2 | 3,600 | 900 | 1,500 |
| Siblings | 1200 | 22 | 26400 | Siblings | 1 | 16,320 | 4,080 | 0 |
|  |  |  |  |  | 2 | 3,600 | 900 | 1,500 |
| Half-siblings | 1200 | 22 | 26400 | Second-degree | 1 | 16,320 | 4,080 | 0 |
|  |  |  |  |  | 2 | 3,600 | 900 | 1,500 |
| Avuncular | 1200 | 22 | 26400 |  | 1 | 16,320 | 4,080 | 0 |
|  |  |  |  |  | 2 | 3,600 | 900 | 1,500 |
| Grandparent-grandchild | 1200 | 22 | 26400 |  | 1 | 16,320 | 4,080 | 0 |
|  |  |  |  |  | 2 | 3,600 | 900 | 1,500 |
| First cousins | 1200 | 22 | 26400 | Third-degree | 1 | 16,320 | 4,080 | 0 |
|  |  |  |  |  | 2 | 3,600 | 900 | 1,500 |
| Great avuncular | 1216 | 22 | 26752 |  | 1 | 16,538 | 4,134 | 0 |
|  |  |  |  |  | 2 | 3,648 | 912 | 1,520 |
| Great-grandparent-great-grandchild | 1200 | 22 | 26400 |  | 1 | 16,320 | 4,080 | 0 |
|  |  |  |  |  | 2 | 3,600 | 900 | 1,500 |
| Unrelated | 3000 | 22 | 66000 | Unrelated | 1 | 40,800 | 10,200 | 0 |
|  |  |  |  |  | 2 | 9,000 | 2,250 | 3,750 |
| Total |  |  |  |  |  | 230,242 | 57,561 | 17,338 |

Additionally, we employed the “--keep_phase, --founder_ids, --fam, and --miss_rate 0 --X X” parameters while running *Ped-sim*, along with a def file containing information on pedigree architecture specified by the “--d” option. We created def files containing all combinations of sexes within a specified pedigree and generated all possible combinations of relatedness dyads using a custom bash script. Unrelated pairs are parents from the parent-offspring pedigrees and randomly chosen pairs from separate sibling pedigrees. We ensured that the individuals in the unrelated pairs descended from different founder individuals. Identical pairs were simply randomly selected pairs of genomic data belonging to the same individual (these pairs will differ due to pseudohaploidisation in the next step). Note that the dataset was limited to the 200,000 SNP positions randomly chosen at the final step of founder simulation.

### Postprocessing genotype data and pseudohaploidization

Among the 200K SNPs chosen, we filtered out those with a minor allele frequency (MAF) <0.05 in the founder population to reduce noise in the relatedness estimates using *PLINK* (*32*) (v1.90p) with the “--maf 0.05” option on the founder VCF files. Additionally, we restricted the positions to autosomal SNPs using the “--autosome” option. The subsequent analyses were conducted with the remaining 96,082 autosomal SNP positions on the VCF files produced with the “--recode vcf” option. Then, we randomly haploidised (pseudohaploidised) the genotypes with a custom *Python* script (*33*) representing the low-coverage setting. We also prepared downsampled versions for each pair’s genomes to enhance our dataset and cover various scenarios of overlapping SNP numbers between pairs. We randomly downsampled the overlapping SNP positions to 1,000, 1,500, 2,000, 2,500, 3,000, 3,500, 4,000, 4,500, 5,000, 6,000, 7,000, 8,000, 10,000, 12,500, 15,000, 20,000, 25,000, 30,000, 35,000, 40,000, and 50,000 positions. The downstream analyses are conducted on the original and downsampled versions.

### Mismatch rate calculation between pairs

Using this data, we calculated the mismatch rates between each pair in our dataset. The mismatch rate measures the mean allelic discordance between pairs within a given genomic window, quantifying the proportion of discordant SNP alleles (inverse of IBS) and indicating the degree of genetic divergence. We calculated the mismatch rate as the ratio between the number of discordant alleles and the total number of positions in sliding windows using the *adna_tools* (*30*) *Python* package, specifically the “dynamic_asc.py” script.

We used two sliding window configurations. Our motivation here was as follows: On the one hand, information on IBS provided by mismatch rate per SNP is minimal while combining information across multiple SNPs can allow more accurate IBS estimation (*34*); therefore, combining mismatch rates across the whole genome can possibly provide the best single estimate for IBS (*3*). On the other hand, using information in multiple windows across the genome can provide information to distinguish parent-offspring and siblings; in addition, deep learning models might also learn to remove shared noise among pairs due to recombination (**Figure 3**). Based on this reasoning, we used both fine-scale and broad-scale configurations. The first configuration uses a window size of 200 SNPs and a step size of 50 SNPs. The second configuration uses a window size of 500 SNPs and a step size of 500 SNPs (non-overlapping windows). This broad-scale might reduce noise but provide us with fewer windows. Because we did not know a priori which configuration would perform best, we performed all tests with both configurations and compared the results.

To estimate the coefficient of relatedness for pairs, mismatch rates need to be normalised to the expected mismatch rate in the population for unrelated pairs, as performed by *READ* (*13*) and related methods. We thus randomly selected 15 *Ped-sim*-simulated individuals who were unrelated, resulting in 105 unrelated pairs. We again calculated mismatch rates for these unrelated pairs in our two configurations with the “adna_tools” *Python* package’s “dynamic_asc.py” script. Since the number of windows per chromosome and the start and end positions of the windows differ between pairs, the normalisation could not be accomplished window by window. Instead, we performed normalisation per chromosome. For this, we computed the mean value of mismatch rates for all windows on each chromosome for each unrelated pair (Equation 1) with the “msm_baseline.py” script of “adna_tools” with the “--mode mean” option. For each unrelated pair, for each chromosome, we calculated the median of the mismatch rates across windows in that chromosome. This resulted in 22 normalisation values corresponding to each autosome for a pair (Equation 2). Before further analyses, we normalised all mismatch rates we computed for all pairs (Equation 3). We applied the same procedures (from Equation 1 to 3) to pairs with downsampled SNPs. Finally, we transformed all normalised mismatch rates across all windows, chromosomes, and pairs into the coefficient of relatedness (*r*) using Equation 4 (*12*).

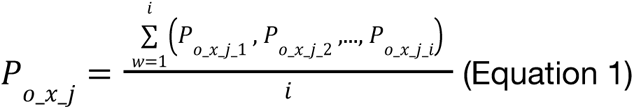

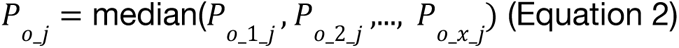

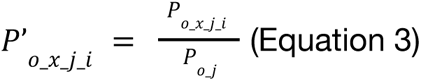

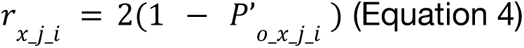

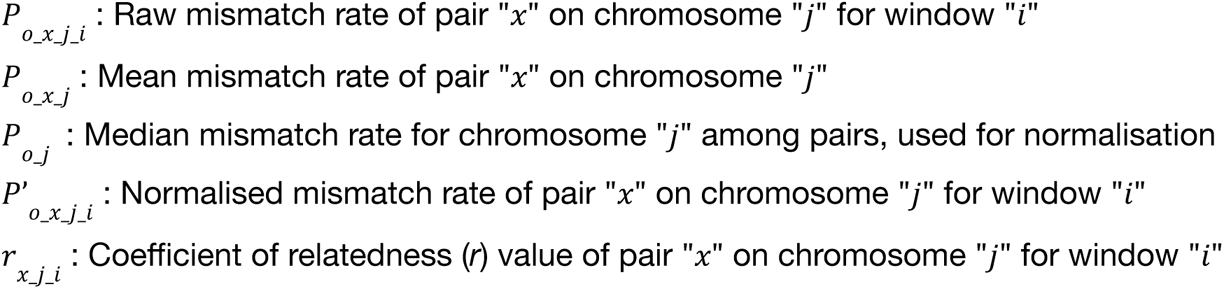

### Preparation of CNN model input

The number of windows, specifically, the sets of *r* value estimates per window for each chromosome of each pair, naturally vary among pairs. CNNs can process inputs of arbitrary lengths; however, the classification layer at the end has to receive a fixed-dimensional vector. Thus, the data has to be mapped to a fixed-dimensional vector at some level of processing. We achieved this through a grid-based summarisation of input data. By applying a predefined grid to the data points, we generated fixed-dimensional matrices for each chromosome. This method ensured consistent input sizes for the downstream deep learning step (see **Figure 2**). We limited the 𝑟 value range from -0.5 to 1.5. While the theoretical range for the *r* is between 0 and 1, we occasionally observed values exceeding one and below 0 for some pairs (see **Figure 3**). We included these outliers in our analysis to retain this information but excluded values beyond -0.5 and 1.5. We counted the number of 𝑟 values within each grid cell from -0.5 to 1.5 and compiled these into fixed-dimensional matrices. Specifically, we divided the range into a 10x10 grid, resulting in 10x10 matrices for each chromosome, which we presumed would be sufficient to capture meaningful information. We chose the size to capture recombination events, known to occur at a rate of 2-4 per human chromosome (*35*). Each matrix represents the distribution of 𝑟 values within that chromosome, maintaining a 2D format. To construct the final input for the deep learning models, we sequentially added the matrices from chromosomes 1 to 22, resulting in a 10x220 matrix. These matrices form the basis of input data for our deep learning models.

**Figure 2:**
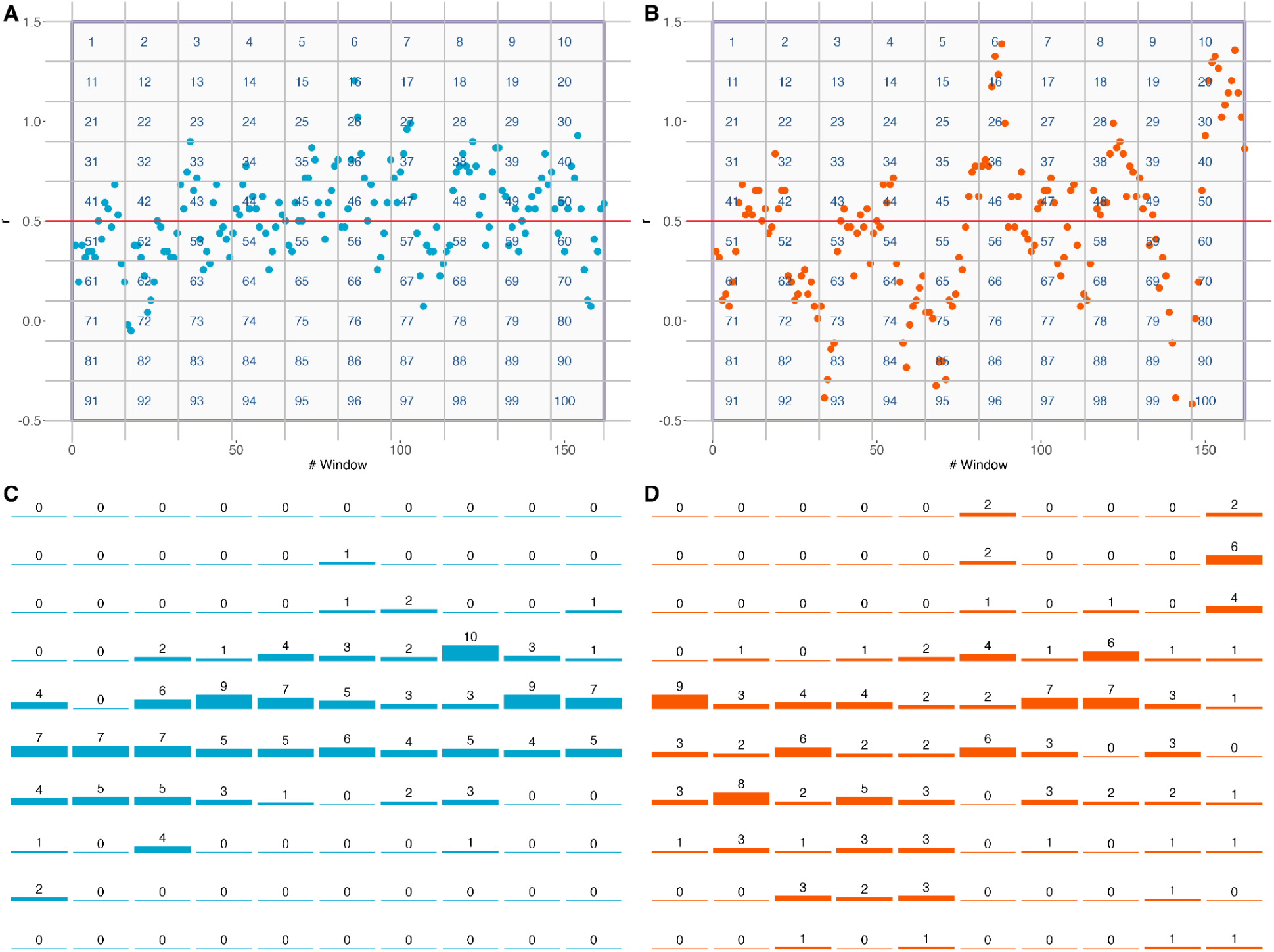
Application of grid template onto *r* value distributions and compilation into fixed-dimensional 2D matrices. Panels **(A)** and **(B)** show the distribution of 𝑟 values across windows of 200 SNPs with a sliding size of 50 SNPs on chromosome 1 for randomly chosen parent-offspring and sibling pairs, respectively. The pairs share exactly 96,082 autosomal SNP positions. Red horizontal lines represent the theoretical expectation for the 𝑟 value for these relatedness categories. The purple square outlines the grid structure, with the number inside each cell corresponding to the matrice position of that grid cell. Panels **(C)** and **(D)** illustrate the count of 𝑟 values in each grid cell position from panels **(A)** and **(B)** for the compiled 2D matrices for parent-offspring and sibling pairs, respectively. The numbers on the top of the bars correspond to the number in that position of the matrix. The higher range of values in the right panel is expected because siblings share parts of their genome in two, single or zero copies.

**Figure 3:**
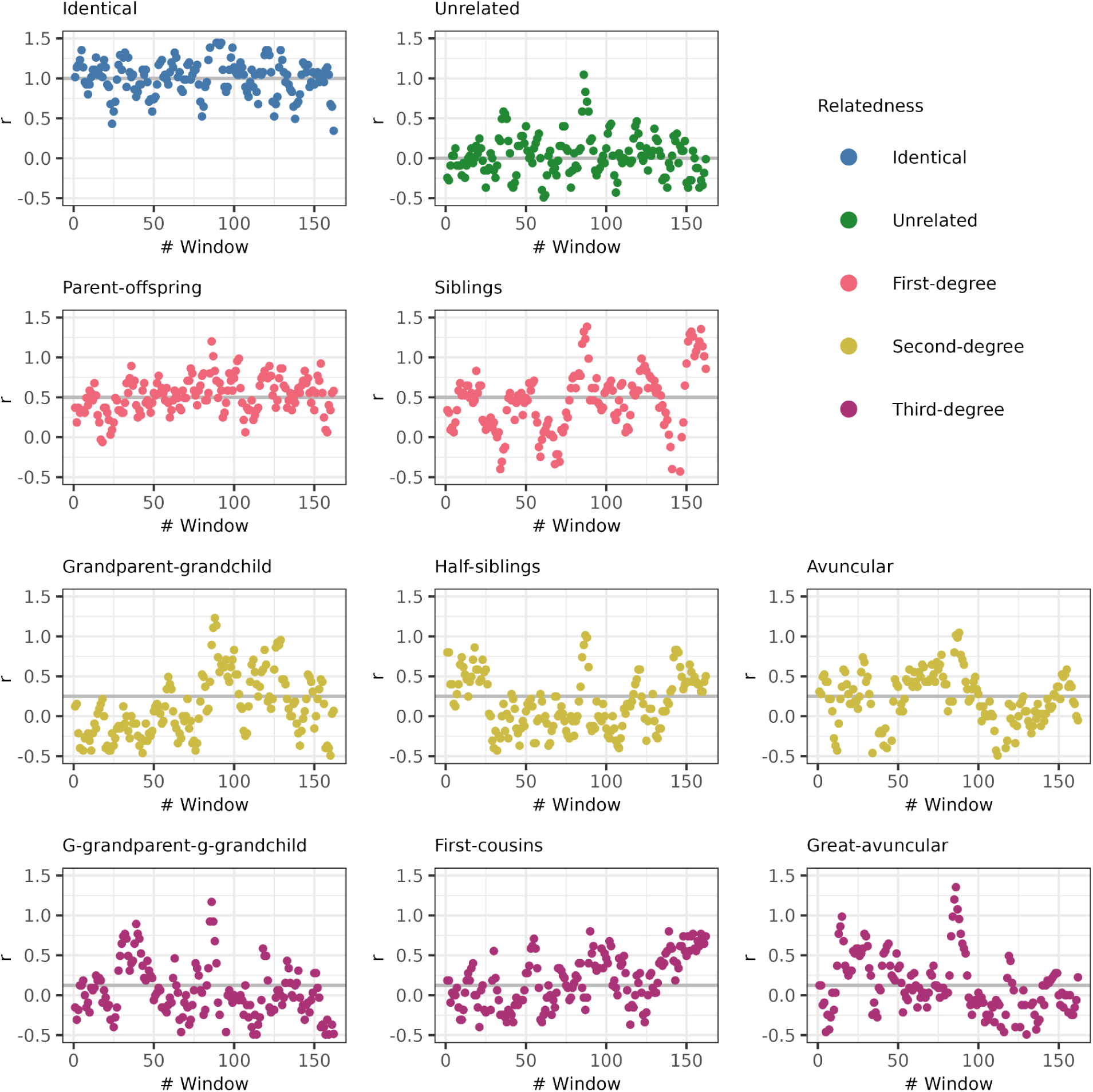
The coefficient of relatedness profiles of each relatedness category simulated in this study. Each panel displays the distribution of coefficient of relatedness 𝑟 values along chromosome 1 for a randomly chosen pair within the specified relatedness category, including identical, unrelated, parent-offspring, sibling, grandparent-grandchild, half-sibling, avuncular, great-grandparent-great-grandchild (denoted “g-grandparent-g-grandchild”), first cousins, and great-avuncular pairs. The 𝑟 values were calculated using sliding windows of 200 SNPs with a step size of 50 SNPs, which is the configuration of Model-A. The pairs share exactly 96,082 autosomal SNP positions. The colours indicate the degree of relatedness. The thick horizontal grey lines represent the theoretical expectation for each specific type of relatedness. **Appendix Figure S4** presents 𝑟 values from the same pairs calculated using 5,000 autosomal SNPs.

## CNN models

### A- Dataset preprocessing

We split the fixed-dimensional matrix dataset into separate training, validation, and test sets (see **Table 1**). We avoided placing different downsampled versions of the same pair into the training, validation, and test sets to prevent data leakage, ensuring the model’s performance evaluation was based on unobserved data. For each 10x220 matrix, we applied label encoding to the target variable, mapping each relationship type label to an integer value. Specifically, all second-degree relatedness types were labelled as "second-degree," and all third-degree relatedness types were labelled as "third-degree" (see **Table 1**).

### B- CNN model architecture

We employed a typical CNN architecture consisting of two main modules: a stack of convolutional layers for feature extraction and fully-connected layers for classification. The architecture included three convolutional layers with kernel sizes of 5, 3, and 3, respectively. The extracted features were flattened and fed into the fully connected layers. To retain positional information in the input, which is crucial for distinguishing between different relatedness categories, we chose not to use pooling layers. The tanh activation function was applied after all layers except the final classification layer, where we used cross-entropy loss for classification, inherently applying the softmax activation function to the final layer to output probabilistic scores.

### C- Training of CNN models

During training, the model adjusted its weights through backpropagation using the Adam optimizer (*36*) from the PyTorch library (*37*). The model’s performance was monitored and validated on held-out validation and test sets. A standard early-stopping method was employed to prevent overfitting, wherein training was halted if the validation loss failed to decrease for a predefined number of epochs (patience). Additionally, we addressed the class imbalance (see **Table 1**) problem by applying class weighting to adjust the training process, where the model is penalised more for misclassifying underrepresented classes. This method helps ensure the model considers all classes fairly, thereby reducing bias and improving overall model performance.

### D- Evaluation and model selection

Since we used two different sliding window configurations to produce our dataset, we trained different models separately on these two configurations. We used various combinations of batch sizes (128 and 256) and learning rates (0.001, 0.0001, 0.00001) for training. The model selection and evaluation process involved three steps. Initially, as a preliminary filtering, the models were evaluated based on their loss curves, focusing on the validation loss. The promising models were then further assessed using the macro average F1 score on the test set (see **Table S1**). The final selection yielded two models for each window size configuration: Model-A, with a learning rate of 0.00001 and a batch size of 256, trained on data generated using sliding windows of 200 SNPs and a step size of 50 SNPs; and Model-B, with the same learning rate but a batch size of 128, designed for sliding windows of 500 SNPs with a step size of 500 SNPs. Both models were used in our final evaluations on simulated and empirical datasets, detailed in **Figures 5B, 5C, 6, and 7**. The metrics reported reflect the models’ capabilities in real-world settings and are distinct from the (synthetic) datasets used for model training and selection.

### Application of *READv2* to the *Ped-Sim* simulated test set

We compared the performance of our models with *READv2* (*3*). We chose this software for three reasons: i) the first version of the algorithm is the most widely used tool in paleogenomic research with ultra-low coverage data (*11*), ii) similar to our approach, it uses average mismatch rates (instead of population allele frequencies) for relatedness estimation, iii) the second version is able to distinguish parent-offspring/sibling pairs and third-degree relatives, unlike the first version. We applied *READv2* to all test set samples to compare its performance with those of our models. We did not apply *KIN* (*9*), another tool that distinguishes parent-offspring and sibling pairs, since it requires mapped genomes in BAM format instead of genotype data. Note that in a recent benchmarking study we showed that *KIN* and *READ* have comparable performances (*11*).

We ran *READv2* on test set samples in batches, allowing *READv2* to calculate normalisation values from each batch. As we created two sets of datasets for Model-A and Model-B, we ran *READv2* on both, resulting in two test results: READv2-A and READv2-B.

### Development of *DeepKin* as a *Python* package

We prepared a *Python* package, *DeepKin*, to process genomic data and predict relatedness categories between sample pairs using the two CNN models, Model-A and Model-B, that we trained with *Ped-sim*-simulated data. The end-user can choose a relevant model, with Model-A as the default. The tool’s initial step involves reading and processing *PLINK*’s (*32*) PED and MAP files to identify individual data and chromosome ranges. Next, it calculates mismatch rates between pairs of individuals across sliding windows on each chromosome. The mismatch data is then aggregated and normalised on a chromosome basis, either using the median mismatch value among all pairs or a user-defined value. Note that we use the median instead of the mean mismatch value among pairs (as in Equation 2) because the median will be robust to the possible presence of related pairs in an empirical dataset (*13*), whereas, in the training data, we chose pairs that were known to be unrelated. The estimated *r* values across the window per chromosome per pair are then compiled into fixed-dimensional matrices. These matrices are converted into a format suitable for CNN, ensuring constant input sizes for the deep learning model. The CNN model is then used to predict the relatedness category for each pair based on the processed data. Finally, the tool generates a comprehensive output file that includes pair names, SNP counts, predicted labels, and prediction probabilities for each class.

### Application to empirical data

We applied our method to published ancient DNA data from two different sites. The first dataset derives from the site of Gurgy ‘les Noisats’ (France), dating back to 4850–4500 BC (*25*). The authors of this study constructed two pedigrees with the genomes they had produced. Among these genomes, we selected 10 individuals unrelated to everyone else and 12 related individuals from the Pedigree B of the study. We excluded fourth-degree related pairs since our method only predicted relatedness up to the third-degree. In total, we analysed 9 parent-offspring pairs, 5 sibling pairs, 21 second-degree pairs, 14 third-degree pairs, and 45 unrelated pairs (see **Appendix Figure S1, Table S2**). The dataset derives from the Rákóczifalva cemetery (Hungary), dating to AD 570-950 (*38*). We selected individuals from the 6th and 8th Rákóczifalva pedigrees, treating pairs between these distinct pedigrees as unrelated. Again, we excluded pairs with relatedness beyond the third degree. In total, we analysed 10 parent-offspring pairs, 3 sibling pairs, 12 second-degree pairs, 5 third-degree pairs, and 49 unrelated pairs (see **Appendix Figure S1**, **Table S2**).

Since our analysis is conducted on each pair separately, we did not calculate the normalisation value only from the analysed pairs. Instead, we calculated it using all individuals from two sites: 94 from the site Gurgy ‘les Noisats’ and 440 from the Rákóczifalva cemetery separately for the relevant dataset. For both datasets, we excluded SNPs with MAF<0.05 using *PLINK*’s (v2.00a4.5) “--maf 0.05” option and restricted the analysis to autosomes using the “--chr1-22” option. To systematically compare our method with *READv2* (*3*), we controlled the number of overlapping SNPs between each pair. After subsetting the overlapping SNPs for each pair, we downsampled the data to 300,000, 200,000, 100,000, 70,000, 50,000, 25,000, 10,000, 5,000, and 3,000 overlapping SNPs using a custom bash script. Some pairs with incompletely covered genomes did not have all downsampled versions due to insufficient numbers of overlapping SNPs, and we only used pairs with exactly the specified number of overlapping SNPs (**Table S2**). We then applied both of our CNN models, Model-A and Model-B, to these pairs and compared the results systematically.

### Calculation of macro average F1 scores

To evaluate the performance of our models and compare them with *READv2*, we calculated the macro average F1 scores for each confusion matrix derived from the test set and empirical paleogenome results. First, we calculated the precision and recall with the formulas 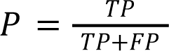 where 𝑇𝑃 is the number of true positives, and 𝐹𝑃 is the number of false positives, and 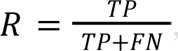 where 𝐹𝑁 is the number of false negatives, respectively. The F1 score was then calculated 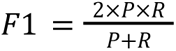 and the macro average F1 score was determined as 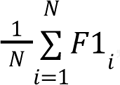 where 𝑁 is the number of classes, and 𝐹1*_i_* is the F1 score for class 𝑖. These scores were calculated for two scenarios; Scenario 1: We combined the parent-offspring and sibling categories into a single "first degree" category, resulting in five categories in the datasets. Scenario 2: We treated parent-offspring and sibling categories separately, resulting in six categories in the datasets. We conducted this analysis in two scenarios because if the pairs share fewer SNPs than a certain threshold, *READv2* does not distinguish between parent-offspring and siblings but simply categorises them as first-degree relationships.

## RESULTS

Our model training using simulated genomes showed that the models could learn relatively fast, with cross-entropy loss reaching stable values on the validation set for both Model-A and Model-B in fewer than 10 epochs (**see Appendix Figure S2**).

### Coefficient of relatedness profiles of *Ped-sim* simulated pairs

Figure 3 presents examples of the distributions of coefficient of relatedness (*r*) values for each relatedness category simulated in this study. These represent the data used to create the training input. Each panel in the figure displays the *r* values along chromosome 1 for a randomly chosen pair from the dataset, ranging from unrelated and identical pairs to different types of first-, second-, and third-degree pairs. The *r* values were calculated using sliding windows of 200 SNPs with a step size of 50 SNPs (the window configuration of Model-A); all the pairs shown share exactly 96,082 autosomal SNP positions. **Appendix Figure S3** presents the same pairs’ *r* value distributions with the Model-B configuration, and **Appendix Figure S4** shows *r* profiles with Model-A using only 5,000 SNPs.

We made five observations. a) The coefficient of relatedness profiles for the simulated pairs generally aligns with theoretical expectations. b) Chromosome-wide variability in *r* is higher for siblings relative to parent-offspring pairs, and variability also increases with genetic distance; these are expected patterns caused by recombination events. c) Part of the intra-chromosomal variation in Figure 3 may be attributed to the presence of the centromere, where low recombination rates appear to create artificial IBD signals. d) The *r* profiles for the same pairs are noisier when pairs share fewer SNPs (**Appendix Figure S4**), highlighting the challenge of accurately predicting relatedness with only a few thousand total SNPs, as the trends visible in Figure 3 tend to fade. e) Finer windows capture more detailed variations in relatedness across the chromosome (see Figure 3), while larger windows provide a smoothed representation, which can obscure detailed patterns but offer a more stable signal (**Appendix Figure S3**).

### The performance of the *DeepKin* models and *READv2* on the simulated test sets

We first studied the relatedness classification performance of *DeepKin* models (Model-A and Model-B) and *READv2* using simulated test set pairs (Figure 4**, Table S3)**. Below, we summarise the results. These show that, overall, the CNN models perform better than *READv2* on this data.

**Figure 4:**
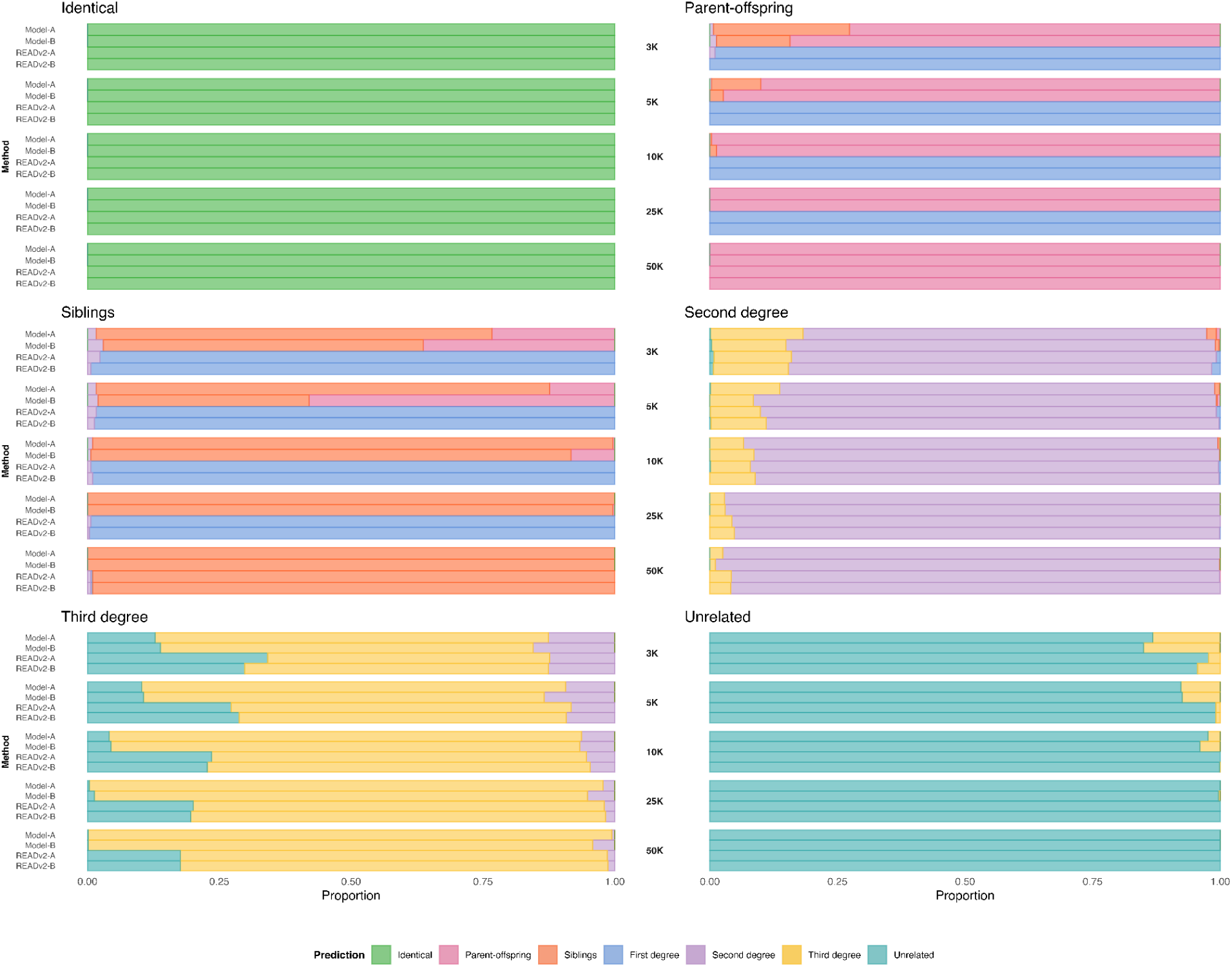
Proportion of predictions across seven different categories of relatedness for the simulated test set. Each bar represents *DeepKin* Model-A, *DeepKin* Model-B and *READv2* with two different setups (READv2-A and READv2-B, the results from two different test sets prepared with different window sizes to be used for comparison with Model-A and Model-B, respectively). The labels in the middle indicate the number of overlapping SNPs between pairs (3K, 5K, 10K, 25K, 50K). The ground truth categories are written in the upper left corner of each panel. The predicted categories by each method are colour-coded, as shown in the legend.

**Figure 5:**
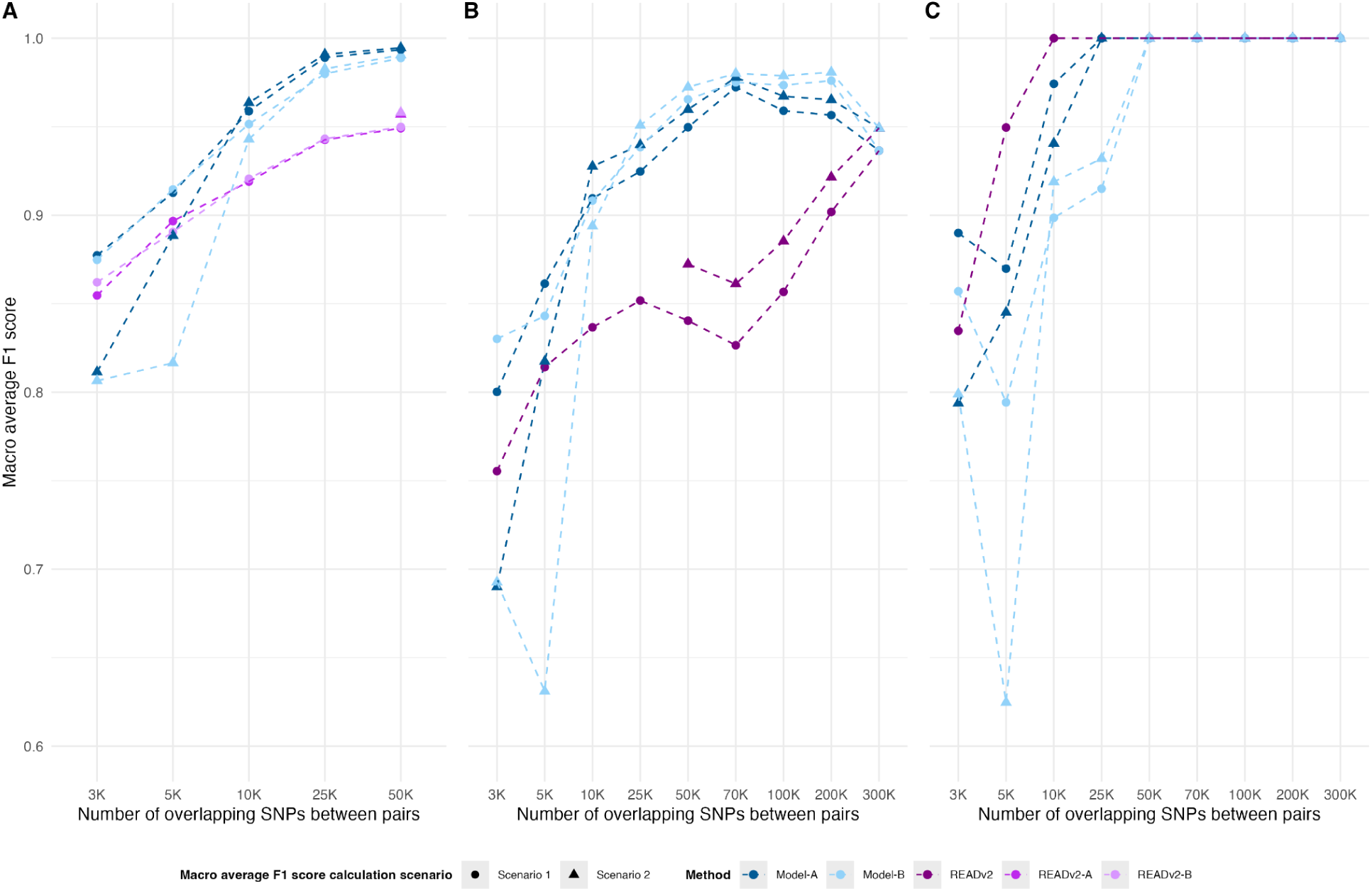
Macro average F1 scores. Panels **(A)**, **(B)**, and **(C)** display the macro average F1 scores of *DeepKin* models and *READv2*, calculated for varying numbers of shared SNPs between pairs. **(A)** depicts scores from the test set. **(B)** depicts scores from the empirical paleogenomic dataset from the Gurgy ’les Noisats’. **(C)** depicts scores from the paleogenomic dataset from the Rákóczifalva cemetery. Scenario 1 describes parent-offspring and sibling categories being combined into a single "first-degree" category when calculating macro average F1 scores, and Scenario 2 describes parent-offspring and sibling categories being treated separately. READv2-A and READv2-B represent the results from two simulated test sets prepared for Model-A and Model-B, respectively (i.e., with different window sizes).

#### Identical pairs

*DeepKin* models and *READv2* consistently achieved perfect classification across all SNP counts.

#### Parent-offspring

Both CNN models demonstrate high accuracy: Model-A has close to 100% at ≥10K SNPs, while Model-B shows slightly lower accuracy. Meanwhile, *READv2* classifies these pairs only at 50K SNPs because it was designed to distinguish between parent-offspring vs. sibling pairs only with relatively high shared SNP counts.

#### Siblings

Model-A correctly classified 75% of siblings even at 3K SNPs, which increased to ≥95% at 10K SNPs and 100% at ≥25K SNPs. Model-B showed slightly lower performance, especially at lower SNP counts. As above, *READv2* classified sibling pairs as siblings only at 50K SNP counts, with 99% accuracy.

#### First degree

When ignoring sibling vs. parent-offspring classification, *DeepKin* models and *READv2* showed similar performances, with ≤3% of sibling pairs misclassified as second-degree at ≤5K SNPs.

#### Second degree

Model-A classified 79% of these pairs correctly at 3K SNPs, improving to 97% at 50K SNPs. Model-B performed slightly better, with 84% accuracy at 3K SNPs and 99% at 50K SNPs. *READv2* had around 83% accuracy at 3K, which improved to 95% at 50K SNPs.

#### Third degree

Model-A correctly classified 75% of such pairs at 3K SNPs, improving to 99% at 50K SNPs. Model-B had slightly lower accuracy, with 71% correctly classified at 3K SNPs and 96% at 50K SNPs. *READv2* performed worst, correctly classifying only around 55% at 3K SNPs and 80% at 50K SNPs.

#### Unrelated

Model-A classified 87% of these pairs correctly at 3K SNPs, improving to 100% at ≥25K SNPs. Model-B also performed well but slightly worse than Model-A, with 85% accuracy at 3K SNPs and 100% at 50K SNPs. *READv2* demonstrated the most robust performance, with around 96% accuracy at 3K SNPs, improving to close to 100% at ≥10K SNPs (see Discussion for an explanation for this difference).

We then used macro average F1 scores to compare the performance of our two CNN models and *READv2* using either 5 or 6 categories, where 5 categories combine parent-offspring and siblings as first-degree relatives (Scenario 1), while 6 categories treat them separately (Scenario 2). In all methods, performance improved visibly with higher SNP counts **(see** Figure 5**, Table S4)**. For 6 categories, Model-A achieved F1 scores of 0.81 at 3K SNPs and 0.99 at 50K SNPs. Model-B also showed the same F1 values. Both models showed slightly better scores for 5 categories, with F1 scores of ∼0.87 at 3K SNPs and ∼0.99 at 50K SNPs. Meanwhile, *READv2*’s F1 scores were calculated only for 5 categories, which remained at ∼0.86 with 3K SNP counts and 0.95 at 50K SNPs.

#### *DeepKin* and *READv2* performance on empirical paleogenomes

We evaluated the performance of the two *DeepKin* CNN models and *READv2* on two different datasets: the Gurgy ’les Noisats’ dataset (*25*) and the Rákóczifalva cemetery dataset (*38*), where pairs shared from 3K to 300K SNPs (Figure 6, Figure 7**, Table S2, Table S5**). We then downsampled these datasets to create pairs with specific numbers of shared SNPs for each category. Since we started with varying numbers of pairs with varying numbers of shared SNPs, the number of pairs per category and SNP number also varied.

**Figure 6:**
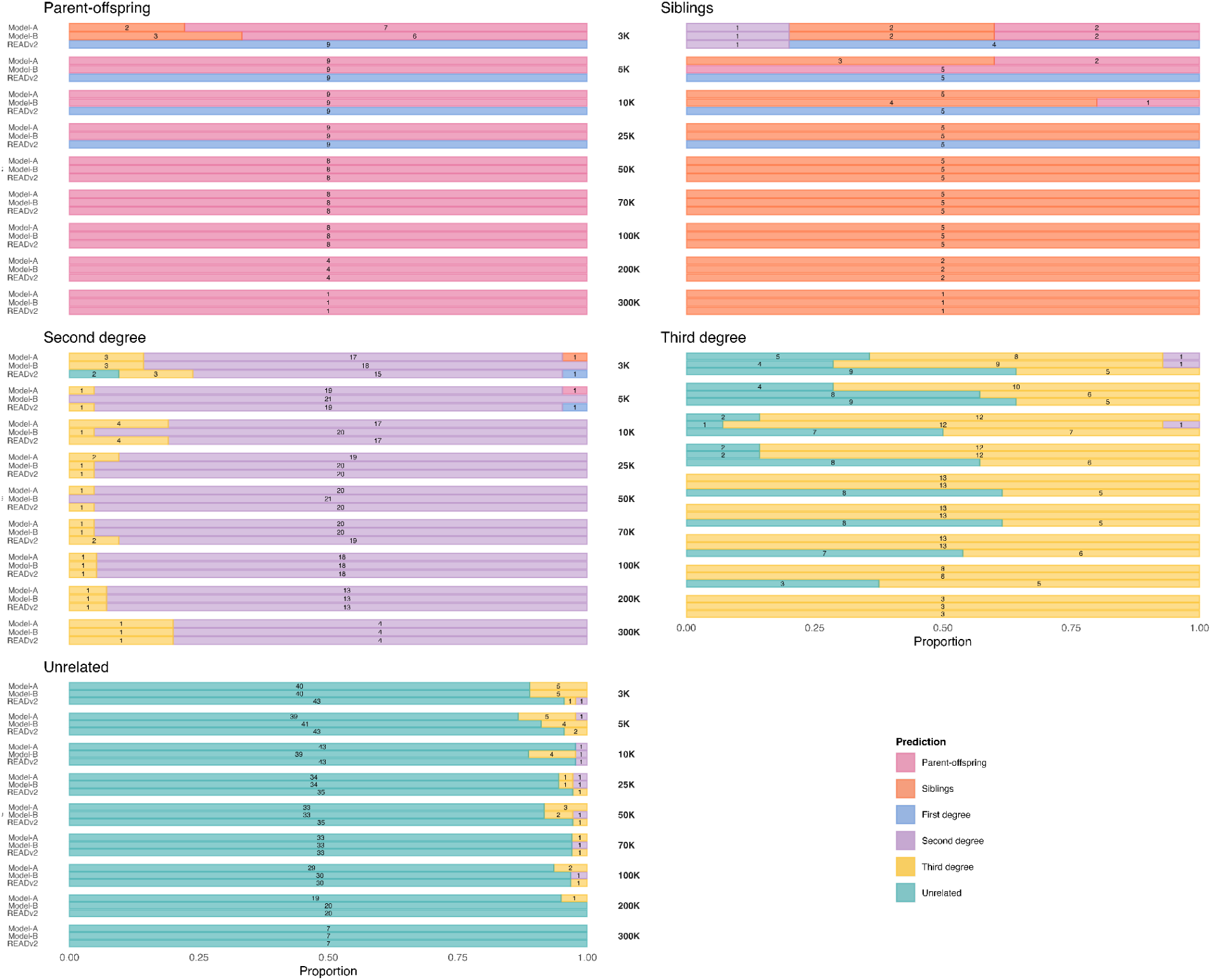
Proportion of predictions across seven different categories of relatedness for the empirical paleogenomes from Gurgy ’les Noisats’. Each bar represents results from a *DeepKin* CNN model or *READv2*. The labels in the middle part indicate the number of overlapping SNPs between pairs (3K, 5K, 10K, 25K, 50K, 70K, 100K, 200K, 300K). The ground truth categories are written in the upper left corner of each panel. The number of pairs in each category is indicated on the bars. The predicted categories by each method are colour-coded, as shown in the legend. The third-degree category did not include any pairs with 300K SNPs.

**Figure 7:**
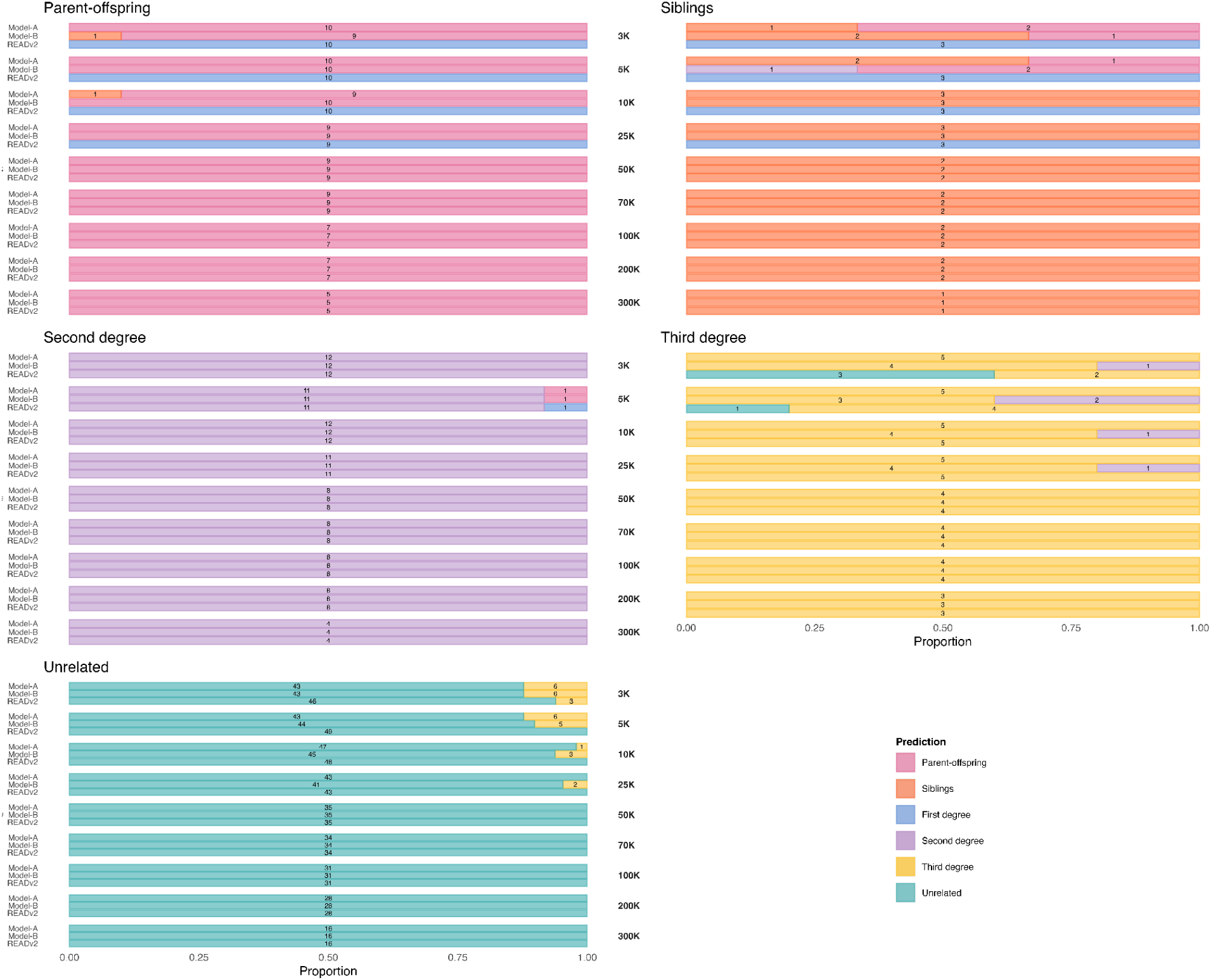
Proportion of predictions across six different categories of relatedness for the empirical paleogenomes from Rákóczifalva cemetery. Each bar represents results from a *DeepKin* CNN model or *READv2*. The labels in the middle indicate the number of overlapping SNPs between pairs (3K, 5K, 10K, 25K, 50K, 70K, 100K, 200K, 300K). The ground truth categories are written in the upper left corner of each panel. The number of pairs in each category is indicated on the bars. The predicted categories by each method are colour-coded, as shown in the legend. The third-degree category did not include any pairs with 300K SNPs.

#### Parent-offspring and siblings

Model-A and Model-B correctly classified all parent-offspring and sibling pairs across the Gurgy and Rákóczifalva datasets at ≥25K SNPs. At ≤10K SNPs, the accuracy of parent-offspring assignment was >80%, but much lower for sibling assignment. Again, *READv2* could only distinguish between parent-offspring and sibling pairs above 50K SNPs. In Scenario 1 (ignoring parent-offspring and sibling distinction), both the *DeepKin* models and *READv2* classified first-degree pairs correctly as first-degree, except for a few mistakes at ≤5K SNPs.

#### Second degree

At 3K SNPs, Model-A and Model-B correctly classified >80% of second-degree pairs in the Gurgy and 100% in the Rákóczifalva datasets. At this SNP count, *READv2* also performed well but with slightly lower accuracy for Gurgy only (71%). Accuracy increased to over 90% at ≥25K SNPs in both datasets using either *DeepKin* models or *READv2*.

#### Third degree

Both datasets had few third-degree pairs (14 and 5). Model-A achieved 8/14 and 5/5 accuracy at 3K SNPs in the Gurgy and the Rákóczifalva datasets, respectively. Model-B performed similarly at 3K SNPs (9/14 and 4/5, respectively). The performance of both models increased with higher numbers of SNPs, reaching 100% at ≥50K SNPs. *READv2* had a lower performance on the Gurgy dataset than the *DeepKin* models (40% at 3K SNPs) but had a similar performance on the Rákóczifalva dataset.

#### Unrelated

*DeepKin* models achieved >88% accuracy at 3K SNPs on the Gurgy and Rákóczifalva datasets, which increased with SNP numbers. *READv2* had >94% accuracy at 3K SNPs in both datasets, outperforming *DeepKin* models.

The macro average F1 scores in these trials highlight differences among the models. Using Scenario 1 (5 categories) on the Gurgy ’les Noisats’ dataset, at 3K SNPs, Model-A achieved an F1 score of 0.80, Model-B scored 0.83, and *READv2* scored 0.76 (Figure 5**, Table S6**). Using Scenario 2 (6 categories) and 50K SNPs, Model-A reached 0.95, Model-B reached 0.97, and *READv2* reached 0.84. Above 50K SNPs, Model-A and Model-B scored 0.95, while *READv2* scored 0.94.

All models performed exceptionally well for the Rákóczifalva cemetery dataset at relatively high SNP counts (Figure 5**, Table S6**). At 3K SNPs, Model-A scored 0.89, Model-B scored 0.86, and *READv2* scored 0.83 for 5 categories. *READv2* performance was higher than *DeepKin* models between 5K-10K SNPs. Meanwhile, Model-A, Model-B and *READv2* all achieved F1 scores of 1.00 at SNP counts of 50K and above for both 5 and 6 categories.

## DISCUSSION

Our study presents *DeepKin*, a novel deep learning framework that we had earlier proposed (*39*), which uses CNNs for accurately identifying relatedness degree when pairs share a low number of SNPs, a challenge in paleogenomics and other fields.

### Training approach

We discuss four choices related to simulations and training. The first choice is training CNNs on normalised summary statistics calculated from SNP data (*r* per window) as input instead of using raw SNP data or the sequence data itself. This choice greatly simplified the training set production. We also hypothesise that it contributed to *DeepKin* being readily generalisable across diverse datasets (see below).

A second choice involved the demographic scenario for genomic data production. As many paleogenomics studies have recently concentrated on studying kinship and social structure in prehistoric West Eurasia (*23–26*), we were motivated to develop models directly applicable to these populations. We therefore chose the Neolithic Anatolian/European demographic model to simulate pedigree founders (see **Methods)**. Note that the Gurgy ’les Noisats’ genetic profiles also belong to the Neolithic West Eurasian pool (*25*), while the Rákóczifalva genomes are distinct, being admixed between East and West Eurasia (*38*). Despite the genetic dissimilarity between the two empirical datasets, our models performed comparably well on the Rákóczifalva genomes. This suggests that our models may generalise effectively across distinct genetic backgrounds. If true, *DeepKin* could be used without training on specific demographic backgrounds. Nevertheless, the effects of more distinct demographic histories on *DeepKin*’s performance, such as bottlenecks, would still be worth investigating in the future.

A third choice was about how realistic the data produced should be. In a previous study, we produced ancient DNA-like sequencing read data, mapped these, and called variants similar to the processing of real paleogenomes, which brought along significant time and computational burdens (*39*). Here, instead, we trained models using simulated variant files (without read simulation, mapping or processing). Nevertheless, our models still performed well on the empirical genomes, indicating robustness.

A final point involved whether to train models using all downsampled versions of the data from 3K to 50K SNPs together or on different SNP counts separately. When pairs share high or low numbers of SNPs, the chromosomal average *r* values remain similar, but the intra-chromosomal variability patterns are altered (Figure 3**; Appendix Figure S4**). Nevertheless, when we compared the effectiveness of model training with the combined downsampled versions versus training separately with data-specific SNP counts, we observed that the former not only maintained classification performance in high-coverage datasets but also enhanced performance on low-coverage datasets **(Table S7)**. This suggests that the information retained from the high-coverage data is transferable to the low-coverage scenarios, easing the latter’s classification task. Also, this simplified the training process without compromising accuracy.

### Window and step size choice

Model-A’s configuration, with a window size of 200 SNPs and a sliding window of 50 SNPs, contrasts with Model-B’s configuration, which uses a window size of 500 SNPs and non-overlapping windows **(Methods)**. Model-A may better capture local variation in the genomic data, such as short IBD segments, which could help distinguish first-degree relative types. Model-B, with its larger and non-overlapping windows, might provide a more averaged view of the genomic data, smoothing out finer details but offering a more stable signal. Model-B’s non-overlapping windows also avoid redundancy, leading to more independent information per data point. When designing the study, we did not have a theoretical expectation which of the configurations might be superior.

In the test set, Model-A overall performed better than Model-B at all SNP counts (Figure 4, Figure 5). In the empirical ancient genome sets, however, Model-B consistently outperformed Model-A in the Gurgy dataset, starting from 25K SNPs (Figure 5, Figure 6). Conversely, in the Rákóczifalva cemetery dataset, both models achieved 100% performance after 50K SNPs, but Model-A was superior at SNP counts lower than 50K (Figure 5, Figure 7). Notably, at the lowest SNP counts Model-A always performed better than Model-B in predicting parent-offspring and sibling pairs, as would be expected.

Since we found no consistent pattern in the performances of the two models across different sets of samples and SNP counts, we did not further explore alternative window configurations. The models may have different strengths depending on the specific dataset and SNP count, and it may be reasonable to use both in parallel.

### Comparison with *READv2*

On the test set, our *DeepKin* models showed consistently superior performance over *READv2* (Figure 4, Figure 5). This can be attributed to the fact that the CNN models were trained using a simulated dataset that closely mirrors the test set’s characteristics. This alignment likely gave *DeepKin* an advantage over *READv2*.

More importantly, comparing *DeepKin* with *READv2* on empirical paleogenomes (with all their complexities), we found that *DeepKin* outperformed *READv2* in nearly all comparisons across various SNP counts and relatedness categories (Figure 5). The only visible exception was the unrelated category: because *READv2* is biased towards underestimating *r* between pairs (similar to other kinship estimation methods (*11*)), it tends to classify all unrelated pairs correctly (when *DeepKin* sometimes assigns these as third-degree). However, this bias also came at a cost to *READv2*, such that third-degree pairs were frequently assigned to the unrelated category. *DeepKin* did not appear to have such an underestimation bias. In summary, the F1 measures on both the simulated data and two empirical datasets suggested that *DeepKin* can be as accurate as *READv2*, if not superior.

### Benefits of CNN models

Using CNNs in *DeepKin* capitalises on their strength in handling grid-like data structures, making them well-suited for genetic data arranged in fixed-dimensional matrices. Unlike traditional methods that treat SNPs or SNP windows independently, our input maintains the genomic context information of per SNP mismatch data, and the CNN models can learn to combine information across genomic segments, improving the detection of specific relationships such as parent-offspring and siblings. CNNs may also be better at learning and removing shared background noise in *r* estimates (Figure 3).

### Data requirements and future directions

Training deep learning algorithms is typically data-intensive, requiring input in the order of millions of samples. Despite our relatively small simulated training set (230K samples), our models still exhibited strong performance. Further, the classification performance on the empirical data was comparable to that of the synthetic test set, even without any fine-tuning. This suggests that our simulated data aligns closely with the empirical samples, at least in the form of *r-*value matrices. This case, where the model has never encountered empirical data during training, could be considered a form of zero-shot learning. Taking into account the high predictive performance obtained, our results highlight the robustness of our approach. However, with more extensive training and validation sets, we anticipate that the models’ performance could further improve, as they would have more data to refine their predictive capabilities and enhance accuracy.

Additionally, one may explore different model hyperparameters, such as the number of convolutional layers, kernel sizes, learning rates, and batch sizes, to further optimise performance. Increasing the number of downsampled versions of the data and expanding the founder population in our simulations could also enhance the model’s robustness and accuracy. These adjustments would provide the models with a richer and more varied dataset, potentially leading to better generalisation and improved predictive performance. Furthermore, different deep learning architectures, especially those specialised for processing sequential data, such as RNNs or Transformers, may be explored as alternative pathways (e.g., using DNA sequences themselves). Nevertheless, we believe that using SNP-based summary statistics as input, as we do here with *DeepKin,* not only greatly simplifies simulations but also facilitates the generalisability of the models.

Another direction for future research is to integrate multiple types of information into the deep learning models by leveraging multi-modal learning techniques. Beyond using mismatch information, as we did here, one could also incorporate genotype likelihoods per SNP (instead of pseudohaploid genotype calls) or population allele frequencies, which could improve the model’s predictive power.

While we trained *DeepKin* for use on ancient genome pairs, the underlying approach has potential applications in other areas of genomics. For instance, the method could be adapted to study relatedness in modern human populations whenever low-coverage or partial genomic data is available. Additionally, the framework could be applied to forensic genetics to identify familial relationships from degraded DNA samples. The approach may also be adopted in conservation genetics in studying wildlife as long as a rough demographic model (or unrelated founder genomes) and genetic maps are available. By extending the application of *DeepKin* beyond paleogenomes, we can leverage its strengths in handling low-coverage data to address various research challenges.

## Acknowledgements

We thank all members of the METU Biological Science CompEvo and of the Hacettepe Human_G groups. We are grateful to METU ImageLab for providing their server to train deep learning models. Special thanks to Gözde Atağ, Igor Mapelli, and Kanat Gürün for their assistance; Wolfgang Haak, Maite Rivollat and Zuzana Hofmanová for sharing data; Pavlos Pavlidis, Aybar Can Acar, Maël Lefeuvre and Céline Bon for their valuable discussions. We acknowledge support from the European Research Council (ERC) Consolidator grant H2020 ‘NEOGENE’ (no:772390 to MS) and H2020-WIDESPREAD-05-2020 TWINNING grant ‘NEOMATRIX’ (no: 952317 to MS), and by the ‘TÜBİTAK-2210/A (The Scientific and Technological Research Council of Türkiye)’ for MNG.

## Author contributions

MNG, AY, BK, EA and MS designed the study. MNG produced the simulated data with the support of KBV. SK and TEÜ produced the preliminary deep learning results. MNG and AY implemented the deep learning models. MNG analysed the data, assisted by AY, BK and NEA. MNG, AY and BK implemented the Python package. MNG and MS wrote the manuscript with contributions from all authors.

## Disclosure and competing interests statement

The authors declare no conflict of interest.

## Data availability

The computer code produced in this study is available in the following GitHub repository: https://github.com/MerveNurGuler/DL-kinship-scripts.git

*DeepKin* is available as a Python package with the following DOI: 10.5281/zenodo.13269720 in the following GitHub repository: https://github.com/MerveNurGuler/DeepKin

## Notes

https://github.com/MerveNurGuler/DL-kinship-scripts.git

https://github.com/MerveNurGuler/DeepKin

### Competing Interest Statement

The authors have declared no competing interest.

