## Appendix for "DeepKin: Predicting relatedness from low-coverage genomes and paleogenomes with convolutional neural networks"

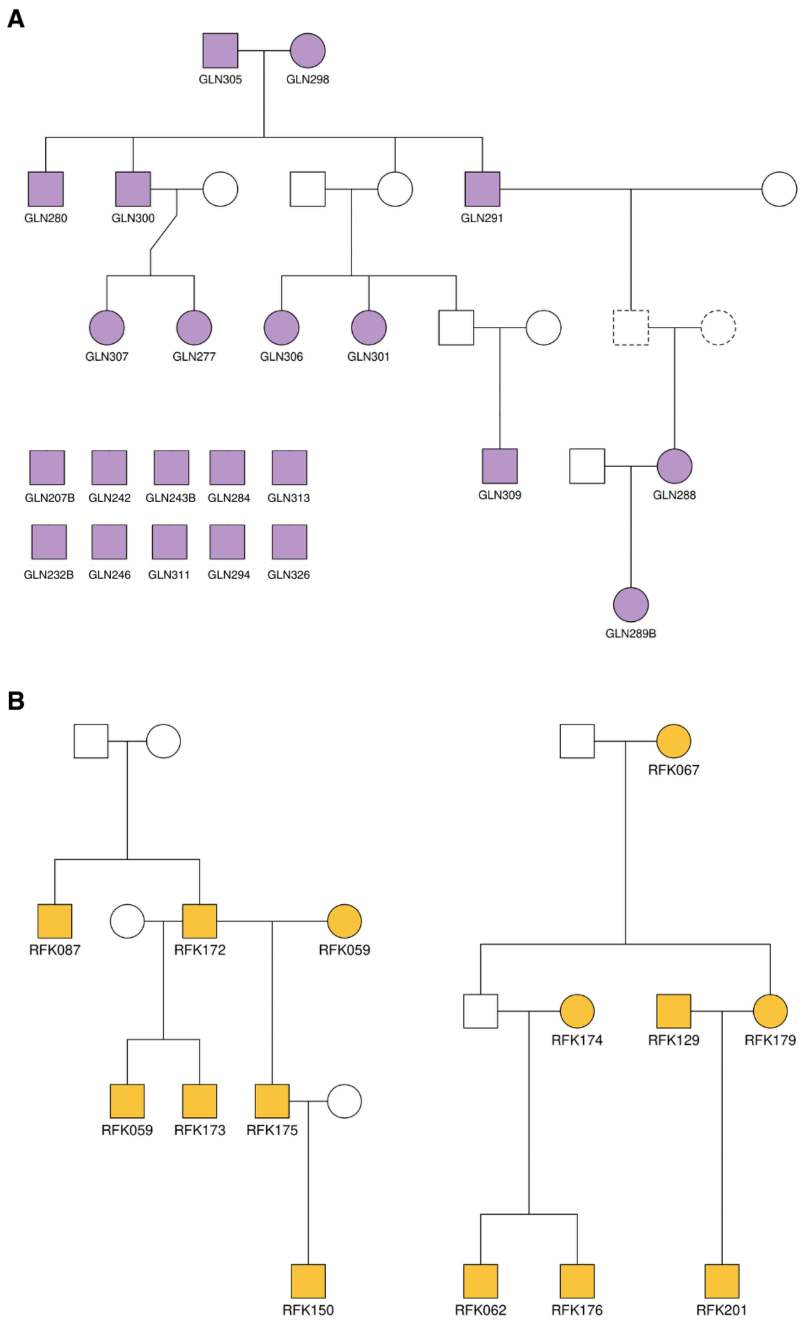


**Appendix Figure S1: Real paleogenome pedigrees used in this study. (A)** Pedigree from the Gurgy 'les Noisats' site in France. This dataset includes 10 unlinked unrelated individuals and 12 related individuals from the study's Pedigree B. **(B)** Pedigree from the Rákóczifalva cemetery in Hungary. Individuals from the 6th (left) and 8th (right) century Rákóczifalva pedigrees were selected, treating pairs between distinct pedigrees as unrelated. Coloured shapes represent individuals included in the analysis, uncoloured shapes represent individuals who were not sequenced, and dashed shapes indicate individuals with uncertain sex.

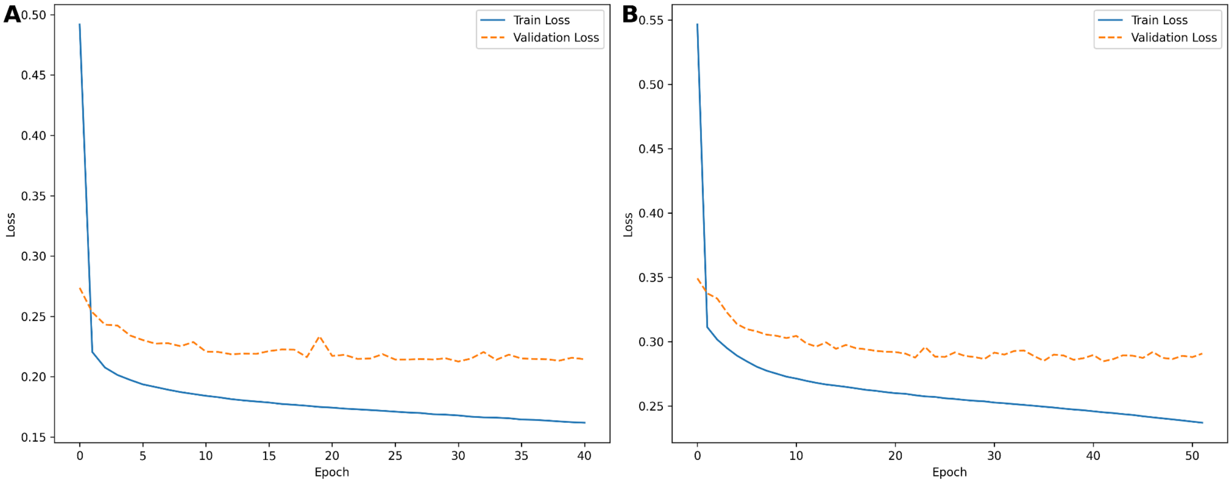


**Appendix Figure S2: Training and validation loss across epochs for two CNN models. (A)** Training and validation loss for the Model A over 41 epochs. **(B)** Training and validation loss for the Model B over 52 epochs. Solid lines represent training loss, while dashed lines represent validation loss.

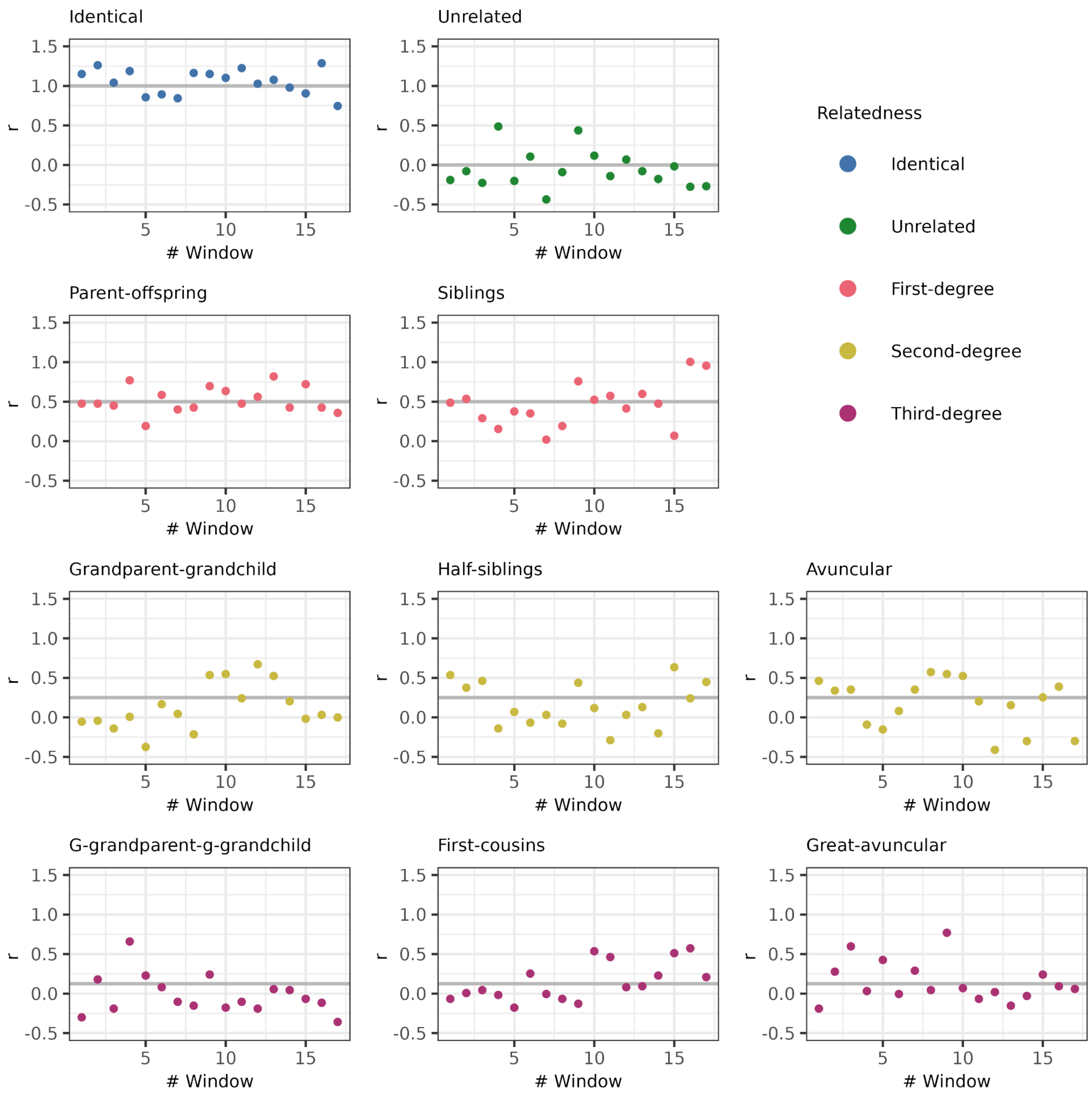


**Appendix Figure S3: The coefficient of relatedness profiles of each relatedness category simulated in this study.** Each panel displays the distribution of coefficient of relatedness $r$ values along chromosome 1 for a randomly chosen pair within the specified relatedness category, including identical, unrelated, parent-offspring, sibling, grandparent-grandchild, half-sibling, avuncular, great-grandparent-great-grandchild (as g-grandparent-g-grandchild), first cousins, and great-avuncular pairs. The $r$ values were calculated using sliding windows of 500 SNPs with a step size of 500 SNPs, which is the configuration of Model-B. The pairs share exactly 96,082 autosomal SNP positions. The colours indicate the degree of relatedness. The horizontal grey lines represent the theoretical expectation for each specific type of relatedness.

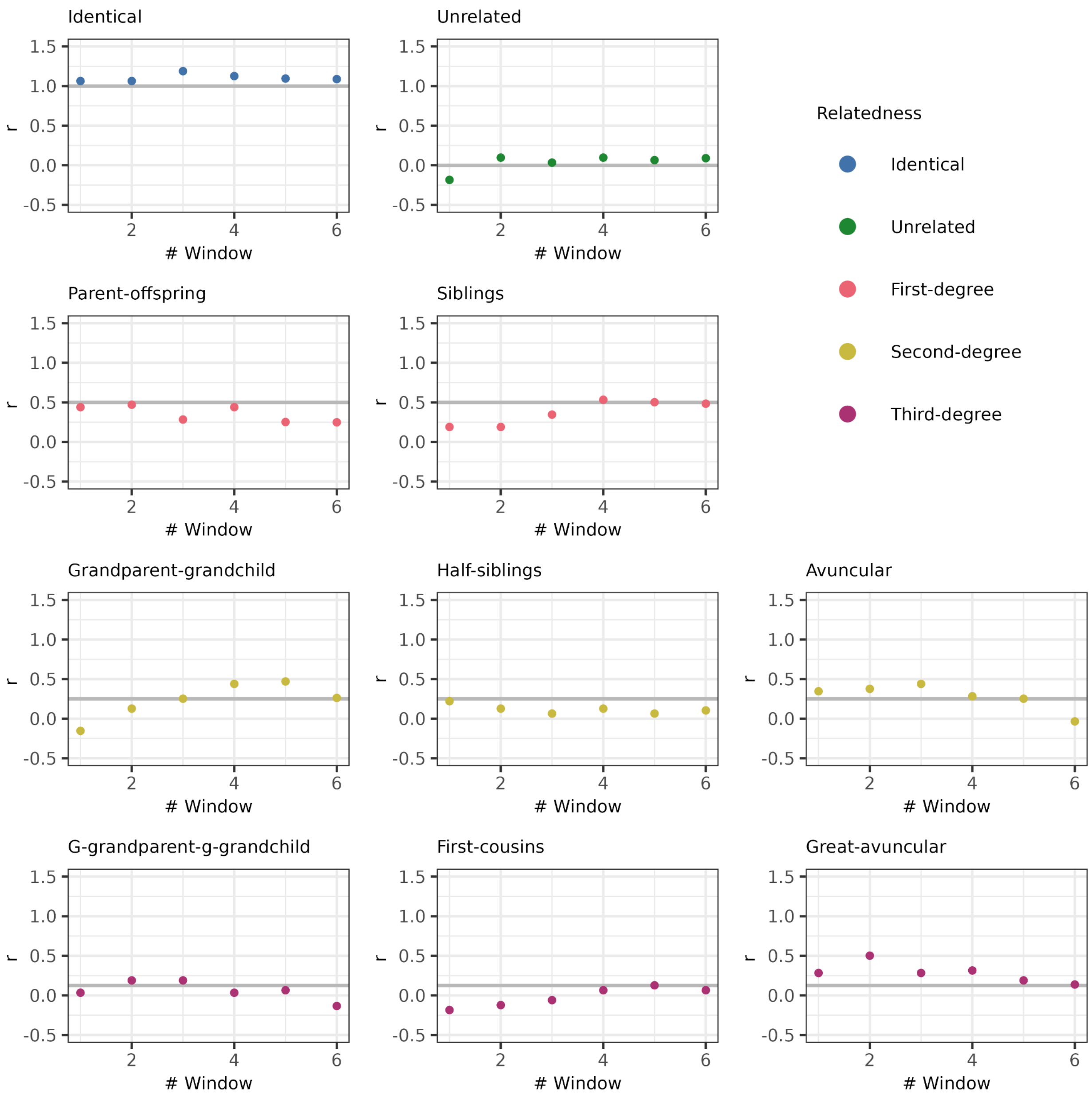


**Appendix Figure S4: The coefficient of relatedness profiles of each relatedness category simulated in this study.** Each panel displays the distribution of coefficient of relatedness $r$ values along chromosome 1 for a randomly chosen pair within the specified relatedness category, including identical, unrelated, parent-offspring, sibling, grandparent-grandchild, half-sibling, avuncular, great-grandparent-great-grandchild (as g-grandparent-g-grandchild), first cousins, and great-avuncular pairs. The $r$ values were calculated using sliding windows of 200 SNPs with a step size of 50 SNPs, which is the configuration of Model-A. The pairs share exactly 5,000 autosomal SNP positions. The colours indicate the degree of relatedness. The horizontal grey lines represent the theoretical expectation for each specific type of relatedness.
